# RNAlign2D – a rapid method for combined RNA structure and sequence-based alignment using a pseudo-amino acid substitution matrix

**DOI:** 10.1101/2020.08.03.233825

**Authors:** Tomasz Woźniak, Małgorzata Sajek, Jadwiga Jaruzelska, Marcin Piotr Sajek

**Affiliations:** Institute of Human Genetics, Polish Academy of Sciences, Strzeszyńska 32, 60-479 Poznań, Poland; Department of Human Molecular Genetics, Institute of Molecular Biology and Biotechnology, Faculty of Biology, Adam Mickiewicz University, Uniwersytetu Poznańskiego 6, 61-614 Poznań, Poland; RNA Bioscience Initiative, University of Colorado School of Medicine, Aurora, Colorado 80045, USA

**Keywords:** RNA, RNA 2D structure, RNA alignment, structure alignment, RNA secondary structure alignment

## Abstract

**Background:** The functions of RNA molecules are mainly determined by their secondary structures. These functions can also be predicted using bioinformatic tools that enable the alignment of multiple RNAs to determine functional domains and/or classify RNA molecules into RNA families. However, the existing multiple RNA alignment tools, which use structural information, are slow in aligning long molecules and/or a large number of molecules. Therefore, a more rapid tool for multiple RNA alignment may improve the classification of known RNAs and help to reveal the functions of newly discovered RNAs.

**Results:** Here, we introduce an extremely fast Python-based tool called RNAlign2D. It converts RNA sequences to pseudo-amino acid sequences, which incorporate structural information, and uses a customizable scoring matrix to align these RNA molecules via the multiple protein sequence alignment tool MUSCLE.

**Conclusions:** RNAlign2D produces accurate RNA alignments in a very short time. The pseudo-amino acid substitution matrix approach utilized in RNAlign2D is applicable for virtually all protein aligners.

## Background

RNA molecules are central players in various cellular processes, including protein biosynthesis and gene expression regulation [1]. These functions are mainly determined by the structures of RNAs (e.g. tRNA, ribozymes), which are often more conserved than RNA sequences [2]. Bioinformatic tools for multiple RNA alignments enable identification of motifs and domains, which are crucial to predict RNA function. Structural information significantly improves alignment quality, as compared to alignments based solely on sequence information. Thus far, secondary structure data (2D structures) are available for > 100,000 RNAs, and the number of RNAs for which the data are available continues to rise [3] in association with the development of high-throughput experimental methods to analyze 2D RNA structures *in vitro* and *in vivo*(for review see [4]).

Several tools to align the structure of RNA molecules have been developed, such as multiple sequence and structure alignment tools, which are usually based on 2D structure prediction algorithms (e.g., TurboFoldII [5] and MAFFT [6], LocARNA [7] and CARNA [8]). LocARNA and CARNA can also use a fixed 2D structure as input. These tools can be divided into three main types. The first entails implementation of the Sankoff algorithm [9], and structure prediction and alignment are performed simultaneously (e.g. LocARNA [7], CARNA [8] or FOLDALIGN [10]). Sankoff algorithm requires O(N^6^) time, where N denotes the length of the compared sequences [9]. Therefore, to reduce complexity, FOLDALIGN uses several heuristics such as the maximum length of the alignment; a maximum difference between any two subsequences being aligned [10]. LocARNA and CARNA use a simplified energy model based on base pair probability matrices to reduce the run-time [7,8]. Additionally, CARNA aligns RNAs with multiple structures per RNA or entire structure ensembles without committing to a single consensus structure. Instead of scoring the alignment of only a subset of the base pairs, it scores the matches of all base pairs in the base pair probability dot plots, which allows aligning of the entire Boltzmann distributed ensemble of structures [8]. In the second group, alignment is based on the sequence and the generated information is used to perform structure prediction (e.g. TurboFold II [5], RNAalifold [11]). The third group entails tools that first predict the structure and then perform the alignment, such as RNAshapes followed by RNAforester [12,13]. However, the tools mentioned can be slow, especially for the analysis of large numbers of long RNA sequences (e.g., 16S rRNA), where specialized tools designed for a particular RNA family may be more suitable (e.g. SSU-ALIGN [14] for 16S rRNA).

To generate alignments of large numbers of long RNA sequences in a short time, we have developed RNAlign2D, a rapid Python tool that aligns multiple RNA molecules based on 2D structure information. It does so by using a pseudo-amino acid substitution matrix, in which RNA sequence and structure are indicated by the use of 1 of 20 characters combined with the protein aligner MUSCLE [15] The idea of using structural information in the sequence alignment was proposed in the early 90’s [16] and was further implemented in STRAL [17]. Our approach represents an alternative solution, dedicated mainly to aligning RNA molecules with known 2D structures, whose number is still growing. RNAlign2D can be applied to perform alignment of either modified or unmodified RNA sequences as well as RNA sequences that contain pseudoknots. Lastly, the RNAlign2D tool can be customized to be compatible with virtually all multiple sequence alignment tools that perform protein alignment.

## Implementation

### General idea

Sequence alignments of RNA are based on aligning four residues: A, C, G, and U. It is possible to use a similar approach to align secondary structures written in dot-bracket format, where ‘.’ represents unpaired nucleotides, ‘(’ and ‘)’ denote paired nucleotides, and other types of brackets are used in the case of pseudoknots [18,19]. To do so, each dot or bracket is converted into a letter arbitrarily assigned to it. In this way, it is possible to align simple secondary structures containing ‘(’, ‘.’, and ‘)’ using 3 letters from the RNA alphabet. To introduce characters describing (first level) pseudoknots ‘[’ and ‘]’, the alphabet has to be extended to at least five letters. One possible solution is to switch from the RNA alphabet to protein alphabet and use protein alignment tools to align the secondary structure of RNA. The protein alphabet consists of 20 letters, therefore other characters like ‘{’, ‘}’ or ‘<’, ‘>’, representing higher-order (nested) pseudoknots [19], can be added. However, higher-order pseudoknots are rather rare. An alternative solution is a combination of RNA secondary structure with its sequence, creating the pseudo-amino acid sequence described below.

### Pseudo-amino acid conversion

As described above, there are two ways to utilize 20 characters of the protein alphabet to represent RNA structure:

1. use dot bracket notation ‘.’, ‘(‘, ‘)’, ‘[‘, and ‘]’ for dot-bracket structures in combination with RNA sequence (20 combinations) to represent each of the RNA nucleotides and the secondary structure assigned to it (e.g., A and ‘.’ when the A nucleotide is in a single-stranded region),
2. arbitrarily assign one of the letters from the protein alphabet to structural elements from dotbracket notation without combining it with RNA sequence.

In this way, it is possible to convert secondary structure or secondary structure with RNA sequence to a new sequence that utilizes the protein alphabet – the pseudo-amino acid sequence. This process is fully reversible, therefore the secondary structure (together with RNA sequence in the first case) can be easily obtained from pseudo-amino acid sequence. However, pseudo-amino acid sequences have nothing to do with the protein sequences encoded in mRNA, except for using the same alphabet.

Both approaches to the conversion have their drawbacks. In the first case, there are limitations for higher-order pseudoknots – they are treated as unpaired regions to keep proper pairing for remaining base pairs. In the second case, there is no information about RNA sequence that may help prepare better alignment.

Details regarding the conversion into all 20 combinations are shown in Figure 1B and Supplementary Figure 1B.

**Figure 1.**
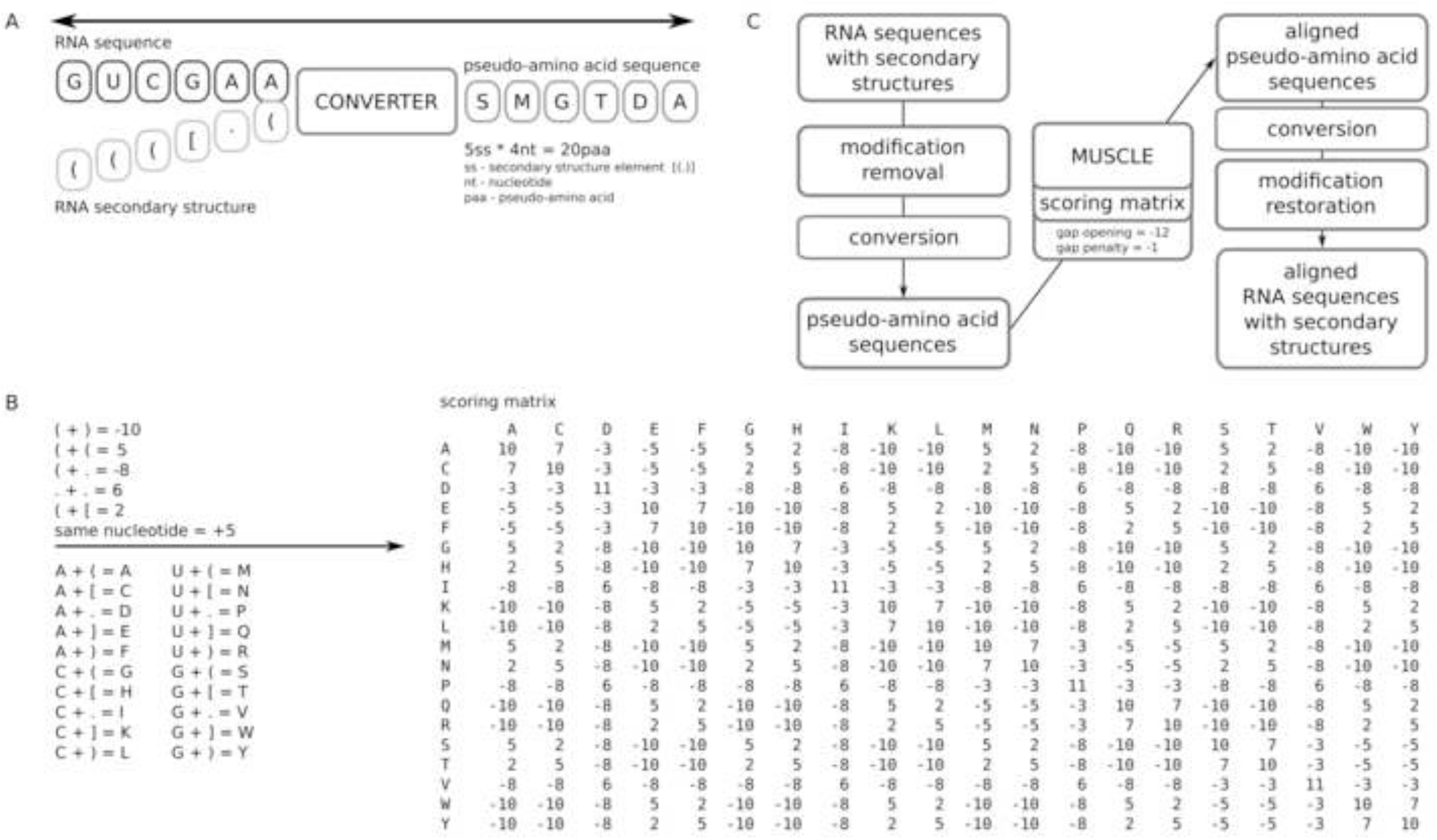
Schematic representation of the RNAlign2D workflow. (A) Basic concept of RNA sequence-structure conversion to a pseudo-amino acid sequence. (B) Conversion of 20 RNA sequence-structure elements to pseudo-amino acids and their scores (left) and the default scoring matrix (right). (C) Block diagram of the RNAlign2D workflow.

It is noteworthy that pseudoknots may be defined in two ways: ((([[[…)))]]] represents exactly the same structure as [[[(((…]]]))). Therefore, we introduced an additional tool that uniformly converts such structures into one common notation.

After the conversion of RNA sequences to pseudo-amino acids, the running of a multiple sequence alignment program dedicated to protein sequences provides the most adequate structural RNA alignment. The MUSCLE program provides such a function for RNAlign2D, utilizing a scoring matrix dedicated to RNA structural alignment. The default scoring matrix for sequence and structure conversion is shown in Figure 1B, and for structure-only conversion, in Supplementary Figure 1B.

### Scoring matrix

Scoring matrix was automatically generated using a selected set of parameters describing scores for pairs of dot-brackets. Different scores are assigned to the same type of bracket or two dots, opposite brackets, different brackets, brackets and dots. Moreover, there is an additional bonus for the same sequence in the aligned molecules. In total, there are eight parameters, including gap opening and gap extension penalty. Theoretically, it is possible to introduce more parameters or even to treat each entry in the matrix separately, but it will most likely lead to overfitting, as there are not enough aligned sequences that can be used to calculate the scoring matrix in this way. To perform an optimal alignment, every parameter of the scoring matrix was optimized using BraliBase 2.1 [20] k7 dataset (further excluded from benchmarks). Optimization lasted 50 iterations and was performed with 18 sets of starting parameters (part of them selected randomly and the rest arbitrary) to reduce risk of local optimum. In each step values in range <current value −4, current value +4> were tested. In case of a higher score, a new value was set, until optimization was complete, in case of equal score there was random chance to change value to the new one. For optimization purposes, SPS score + PPV score + 2 * structural distance score values were used, with maximizing SPS and PPV and minimizing structural distance. Structural distance score values were calculated as 1 - (mean_distance/ length of sequence). The final values for parameters are as follows: same brackets: +5; two dots: +6; different brackets with the same orientation: +2; brackets with different orientation: −10; bracket and dot: −8; bonus for the same sequence: +5; gap opening: −12; gap extension: −1.

### The RNAlign2D tool

RNAlign2D is a command line tool written as a Python3 script that works in UNIX-based operating systems. It is installed via python3-setuptools. Furthermore, MUSCLE aligner requires separate installation. RNAlign2D was tested with MUSCLE v3.8.31. RNAlign2D performs the following processing steps (Figure 1C): (1) removes modifications from RNA sequences (it uses abbreviations for modifications from the MODOMICS database [21]); (2) converts the secondary structures and sequence of the RNAs to pseudo-amino acid sequences; (3) runs the MUSCLE program with the given sequence, scoring matrix, and penalties for gap opening and extension; (4) converts the aligned pseudo-amino acid sequences to RNA sequences and secondary structures; (5) restores the original modifications to each sequence. RNAlign2D consists of an alignment tool, predefined matrices, a scoring matrix creation tool, a modification removal tool, consensus structure calculation tool, and a pseudoknots standardization tool. It also contains a set of files with test sequences to perform alignment.

RNAlign2D can be run by simply writing the following command in a terminal: *rnalign2d -i input_file_name -o output_file_name*. Additional flags allow the users to provide their own scoring matrix, apply penalties for gap opening and/or extension, to choose the running mode (‘simple’ or ‘pseudo’), or to standardize pseudoknot notations. Additionally, the script ‘create_matrix.py’ allows the user to define a customized scoring matrix and calculate_consensus.py to calculate consensus structure for a given alignment. The ‘pseudo’ mode is experimental feature for higher order pseudoknots, where sequence is not taken into account and it should be used sparingly.

The input file used to run RNAlign2D in both ‘simple’ and ‘pseudo’ mode is a FASTA-like file including a header followed by a line containing the sequence and 2D structure in a dot-bracket format. In the ‘pseudo’ mode, the sequence line in this file is omitted during conversion and alignment. When structures with higher-order pseudoknots are analyzed in the ‘simple’ mode, the residues in higher-order pseudoknots are treated as unpaired residues to ensure proper pairing of remaining residues. Moreover, RNAlign2D ‘normalizes’ structures to ensure that pseudoknots are written in a uniform way.

## Results

### Benchmark – sum-of-pair-scores and positive predictive values

RNAlign2D was compared with LocARNA, CARNA, MAFFT, TurboFold II, and STRAL, using BraliBase 2.1 [20] and data from the RNAStralign database [5] as benchmark datasets. LocARNA and CARNA were selected because they can use fixed 2D structure as input. MAFFT and TurboFold II showed the best performance in the previously published benchmark [5]. STRAL utilizes structural information to perform sequence alignment [17]. The sum-of-pair scores (SPSs), positive predictive values (PPVs), structural distance, and running times for each program were calculated.

For alignment of the BraliBase 2.1 benchmark dataset, RNAlign2D, LocARNA, and CARNA generated similar mean SPSs and PPVs for all datasets, which ranged from 0.89 to 0.93 (Figure 2). The mean PPV ranged from 0.71 (k15, LocARNA) to 0.91 (k2, RNAlign2D, LocARNA, and CARNA) (Figure 3). For MAFFT, STRAL, and TurboFold II, those values were lower for most datasets, except PPV for k15, where MAFFT and TurboFold II were comparable to RNAlign2D, LocARNA, and CARNA.

**Figure 2.**
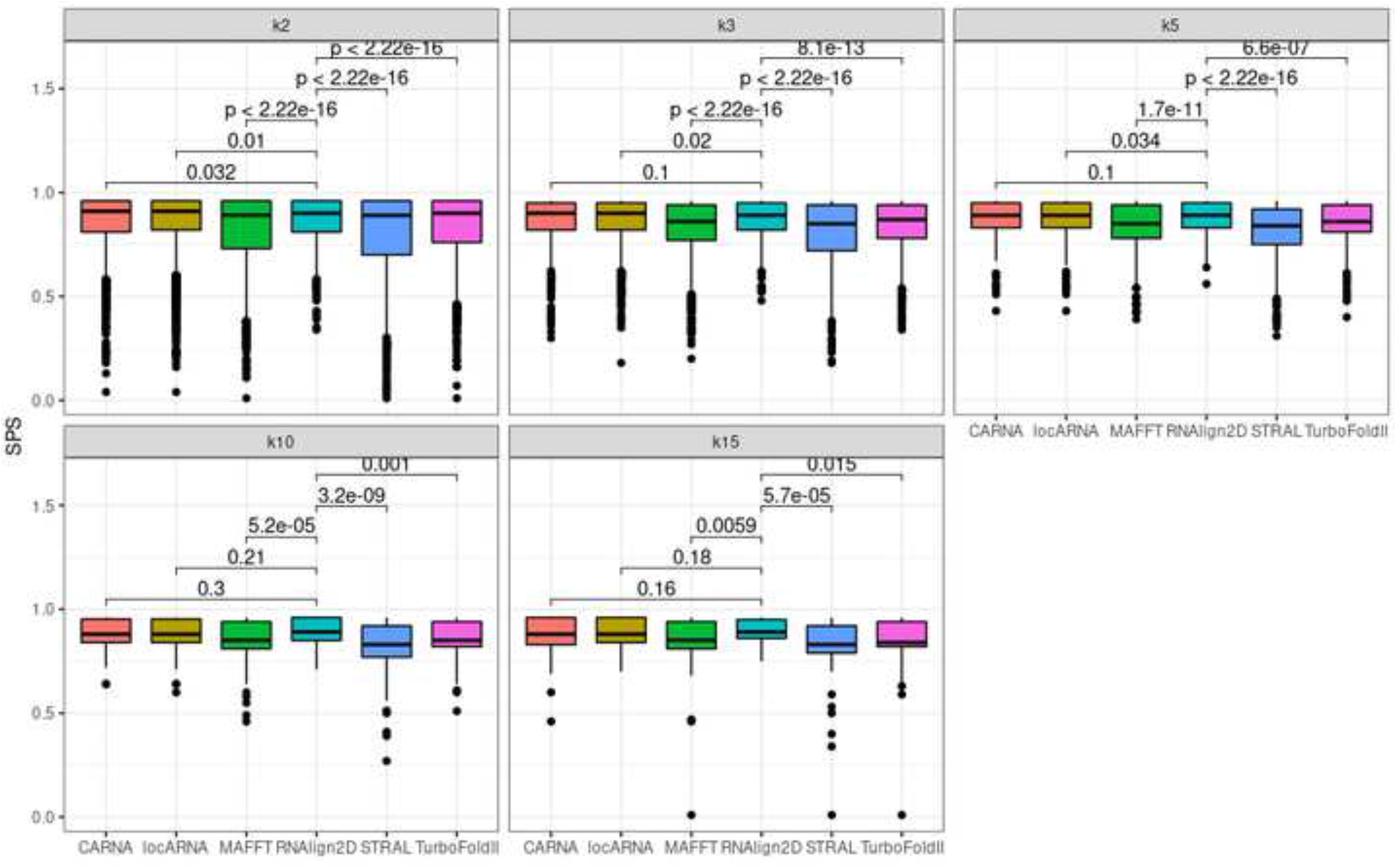
Box and whisker plots comparing sum-of-pair scores (SPSs) generated for the alignment of all sequences in the BraliBase 2.1 benchmark dataset with RNAlign2D, CARNA, LocARNA, MAFFT, STRAL, and TurboFold II (k indicates the number of aligned sequences). P-values were calculated using two-sided t-test.

**Figure 3.**
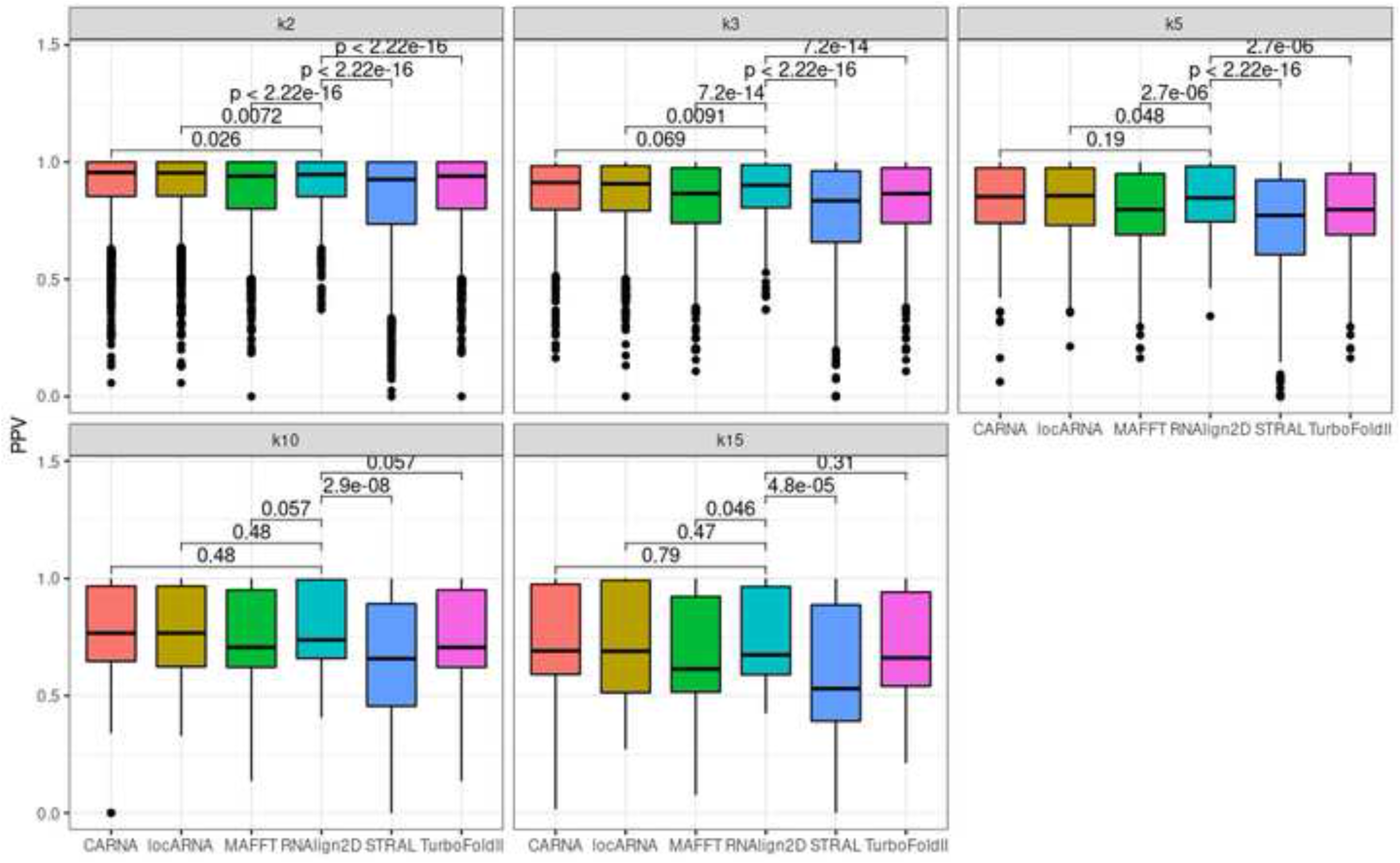
Box and whisker plots comparing positive predictive values (PPVs) generated for the alignment of all sequences in the BraliBase 2.1 benchmark dataset with RNAlign2D, CARNA, LocARNA, MAFFT, STRAL, and TurboFold II (k indicates the number of aligned sequences). P-values were calculated using two-sided t-test.

The RNAlign2D scoring matrix was optimized on the k7 dataset from BraliBase2.1. To ensure that there was no overfitting, we recalculated SPSs and PPVs on the k2, k3, k5, and k10 datasets without alignments containing ≥ 2 (k2, k3), ≥ 3 (k5), and ≥ 5 (k10) common sequences with the k7 dataset for RNAlign2D. We observed only minor, non-significant changes, which means that our scoring matrix is not over-fitted.

To check the performance of alignment of RNA sequences from specific RNA families, we used the RNAStralign benchmark dataset [5]. When this benchmark dataset was aligned, TurboFold II showed the best performance in case of 16S rRNA and ribonuclease P (RNase P) SPS values, where RNAlign2D was only slightly worse and outperformed other programs. RNAlign2D produced the best alignments for RNase P in terms of PPV values and for telomerase dataset (both SPS and PPV). When signal recognition particle (SRP) RNA sequences were aligned, RNAlign2D outperformed only STRAL, produced very similar alignments to MAFFT (in terms of PPV) and worse than other programs used in the benchmark (Figures 4–5). In general, among alignment of all the analyzed RNAs from different families, alignment of the SRP RNA yielded the lowest SPS and PPV. Examples of alignments for each of the above-mentioned families are shown in Figure 6.

**Figure 4.**
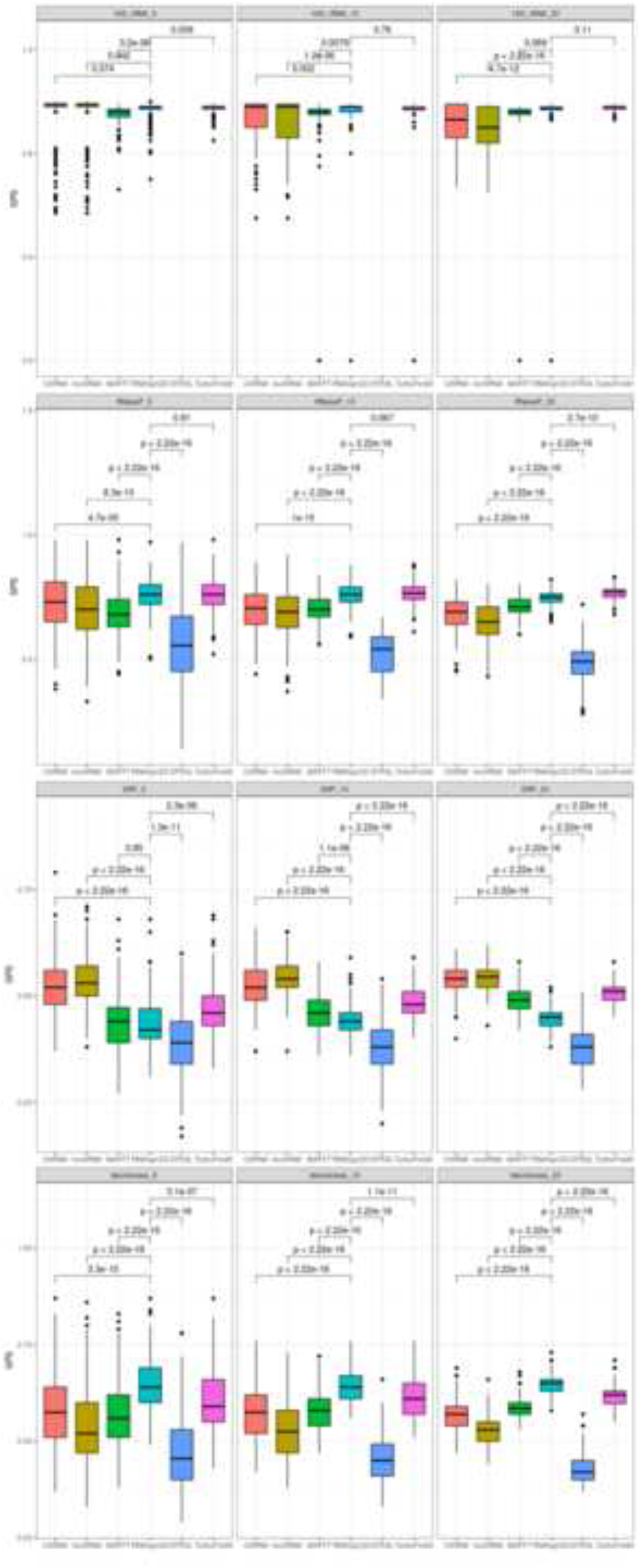
Box and whisker plots comparing sum-of-pair scores (SPSs) for the alignment of 200 groups of 5, 10, and 20 homologous sequences from the entire RNAStralign benchmark dataset with RNAlign2D, CARNA, LocARNA, MAFFT, STRAL, and TurboFold II. P-values were calculated using two-sided t-test.

**Figure 5.**
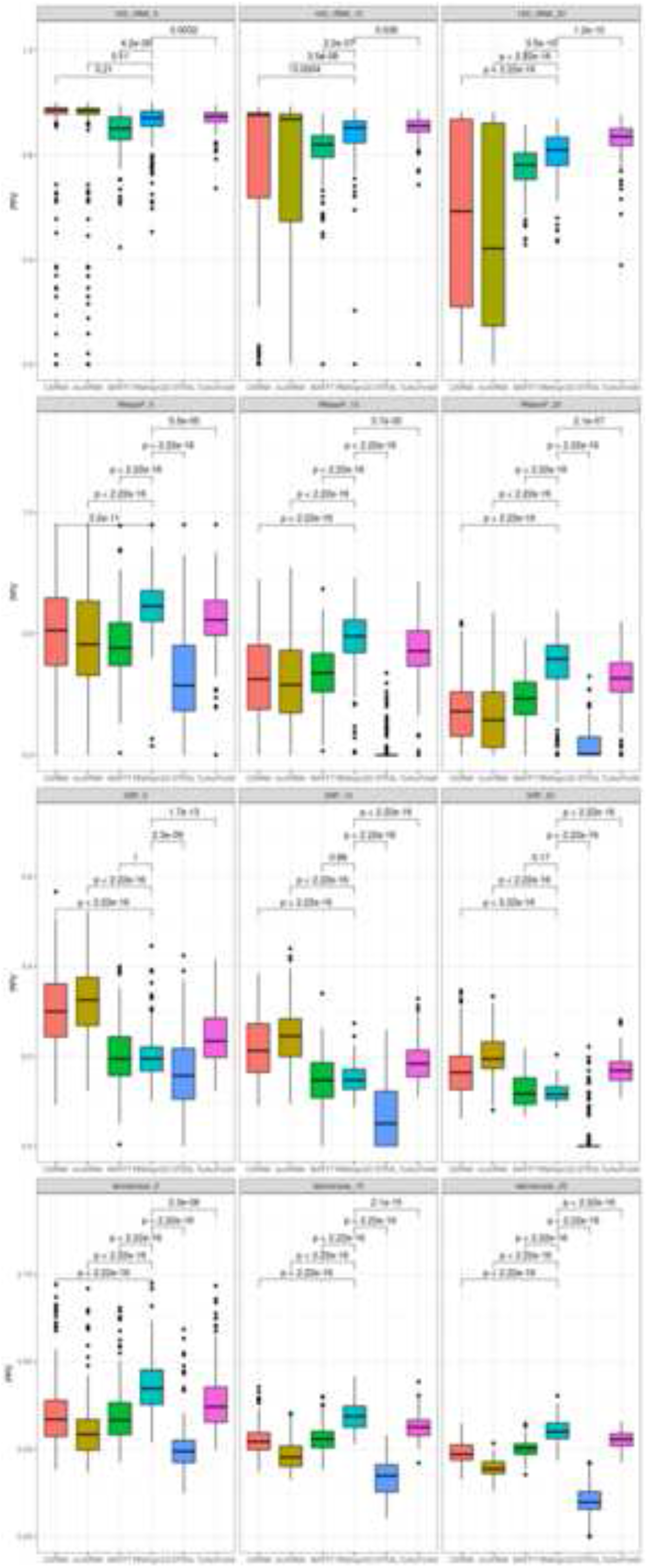
Box and whisker plots comparing positive predictive values (PPVs) for the alignment of 200 groups of 5, 10, and 20 homologous sequences from the entire RNAStralign benchmark dataset with RNAlign2D, CARNA, LocARNA, MAFFT, STRAL, and TurboFold II. P-values were calculated using two-sided t-test.

**Figure 6.**
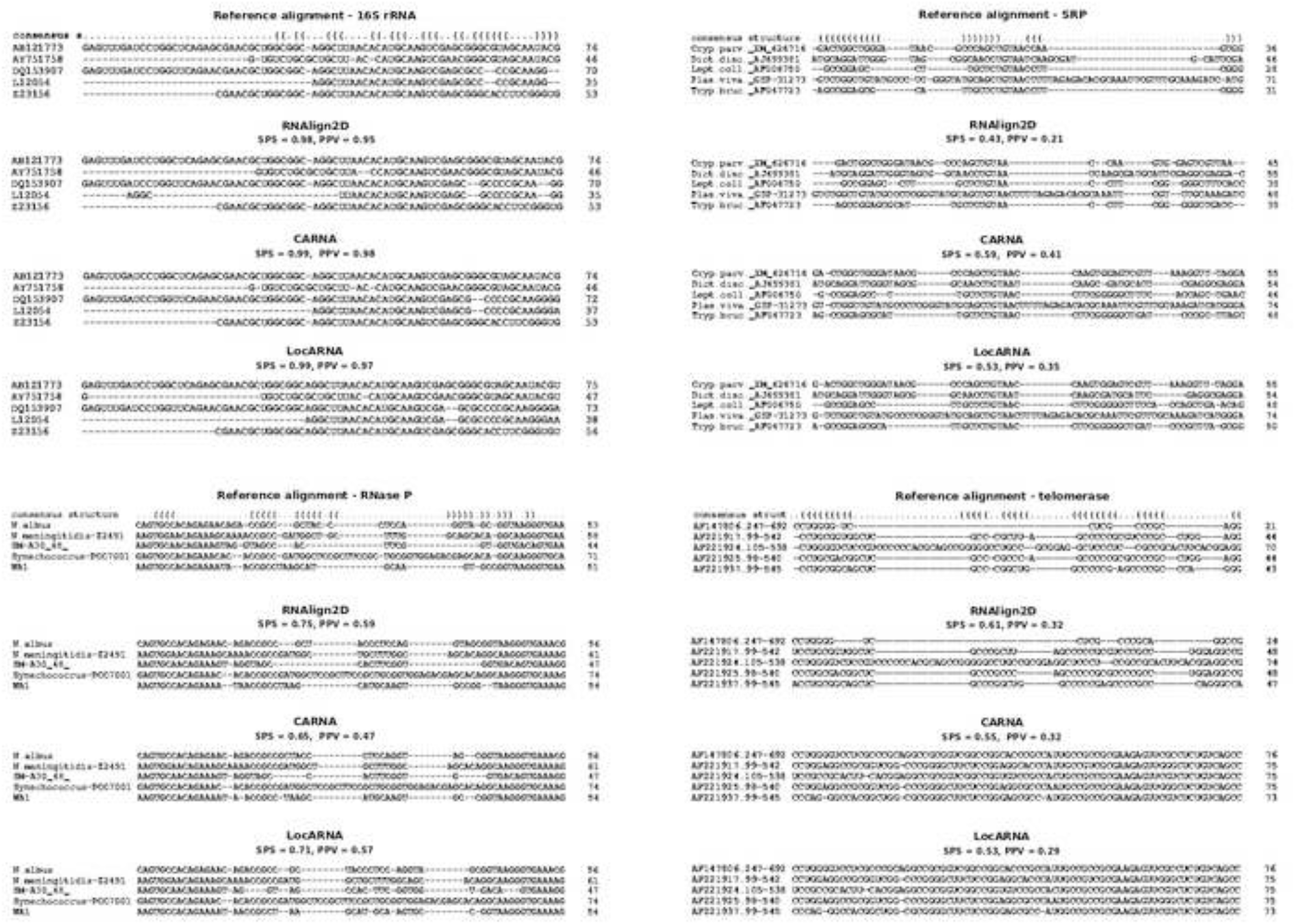
Comparison of alignments produced by tools that utilize known 2D structures for alignment (RNAlign2D, CARNA, and LocARNA) for 16S rRNA, RNase P, SRP, and telomerase families. Examples were chosen from RNAStralign datasets containing 5 sequences. A 75-nucleotide window is shown for each alignment. Numbers on the right side of alignments indicate the length of a particular sequence within the 75-nt window.

The SPSs, PPVs, and standard deviations from the alignment of all datasets with all the alignment tools tested are summarized in Supplementary Table S1.

### Structural distance

As expected, programs that utilize known RNA structures produce better structural alignments than those that predict 2D structures. For the BraliBase2.1 benchmark, RNAlign2D, LocARNA, and CARNA have similar, very low mean structural distances, while for STRAL and TurboFold II these distances are much higher (Figure 7). A similar situation is observed for 16S rRNA and RNase P datasets from the RNAStralign benchmark. For SRP and telomerase datasets, the programs that utilize the Sankoff algorithm outperform RNAlign2D, which in turn outperforms STRAL and TurboFold II (Figure 8).

**Figure 7.**
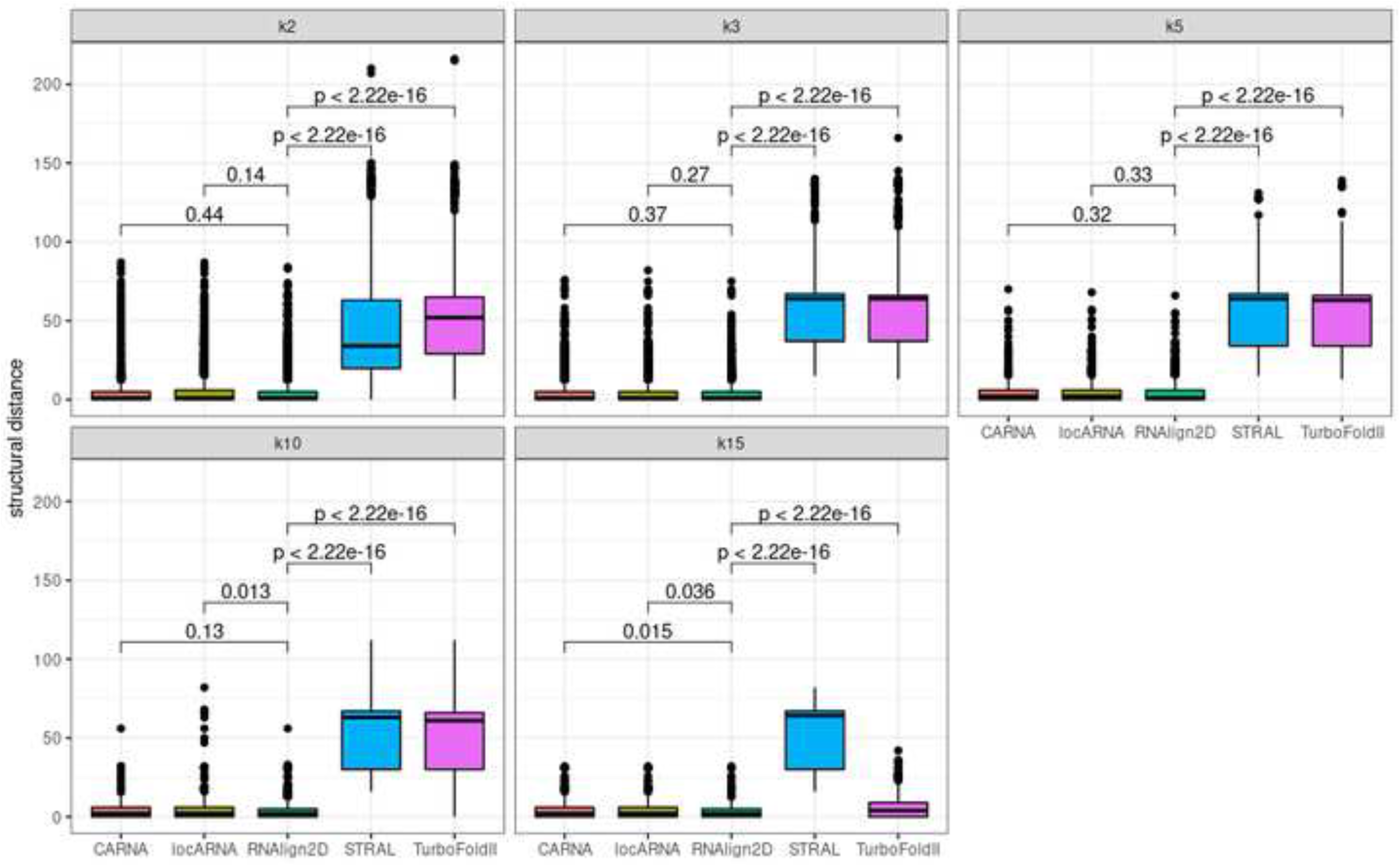
Box and whisker plots comparing structural distances for the alignment of all sequences in the BraliBase 2.1 benchmark dataset with RNAlign2D, CARNA, LocARNA, MAFFT, STRAL, and TurboFold II (k indicates the number of aligned sequences). P-values were calculated using two-sided t-test.

**Figure 8.**
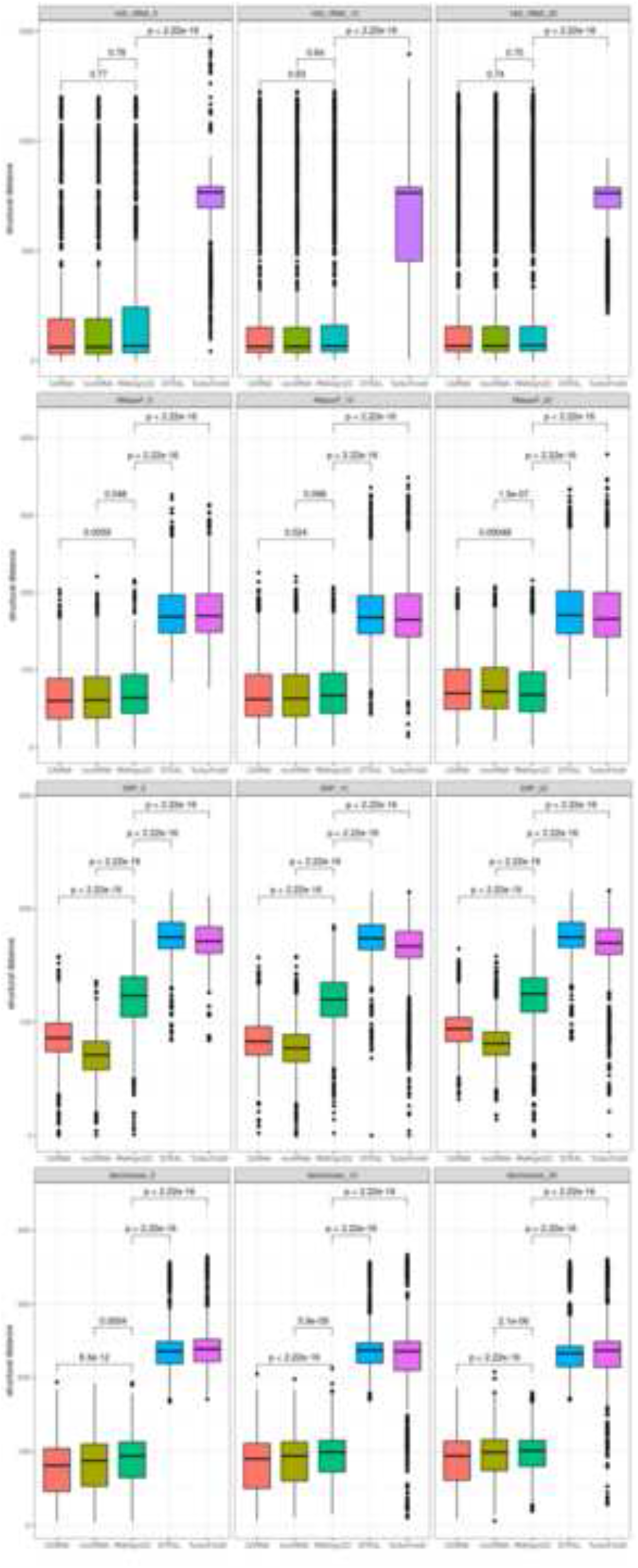
Box and whisker plots comparing structural distances for the alignment of 200 groups of 5, 10, and 20 homologous sequences from the entire RNAStralign benchmark dataset with RNAlign2D, CARNA, LocARNA, MAFFT, STRAL, and TurboFold II. P-values were calculated using two-sided t-test.

### Alignment time

Alignment times from each of the analyzed groups of RNAs from the RNAStralign benchmark datasets were determined and compared. RNAlign2D was the fastest tool for the alignment of datasets containing 20 and 10 molecules (Figure 9), with the alignment time varying from < 1 to 4 s. STRAL had a similar runtime for datasets containing five molecules. However, in the case of 16S rRNA, we were unable to perform alignment with STRAL due to ‘Segmentation fault’ error. Alignment lasted 5–3061 s for LocARNA, 3–34198 s for CARNA, 1–284 s for MAFFT, 24–27252 s for TurboFold II, and between <1 and 20 s for STRAL. Therefore, by simplifying the sequence and 2D structure to pseudo-amino acid sequence as well as using MUSCLE protein aligner, we shortened the alignment time enormously. The obtained results are summarized in Supplementary Table S2.

**Figure 9.**
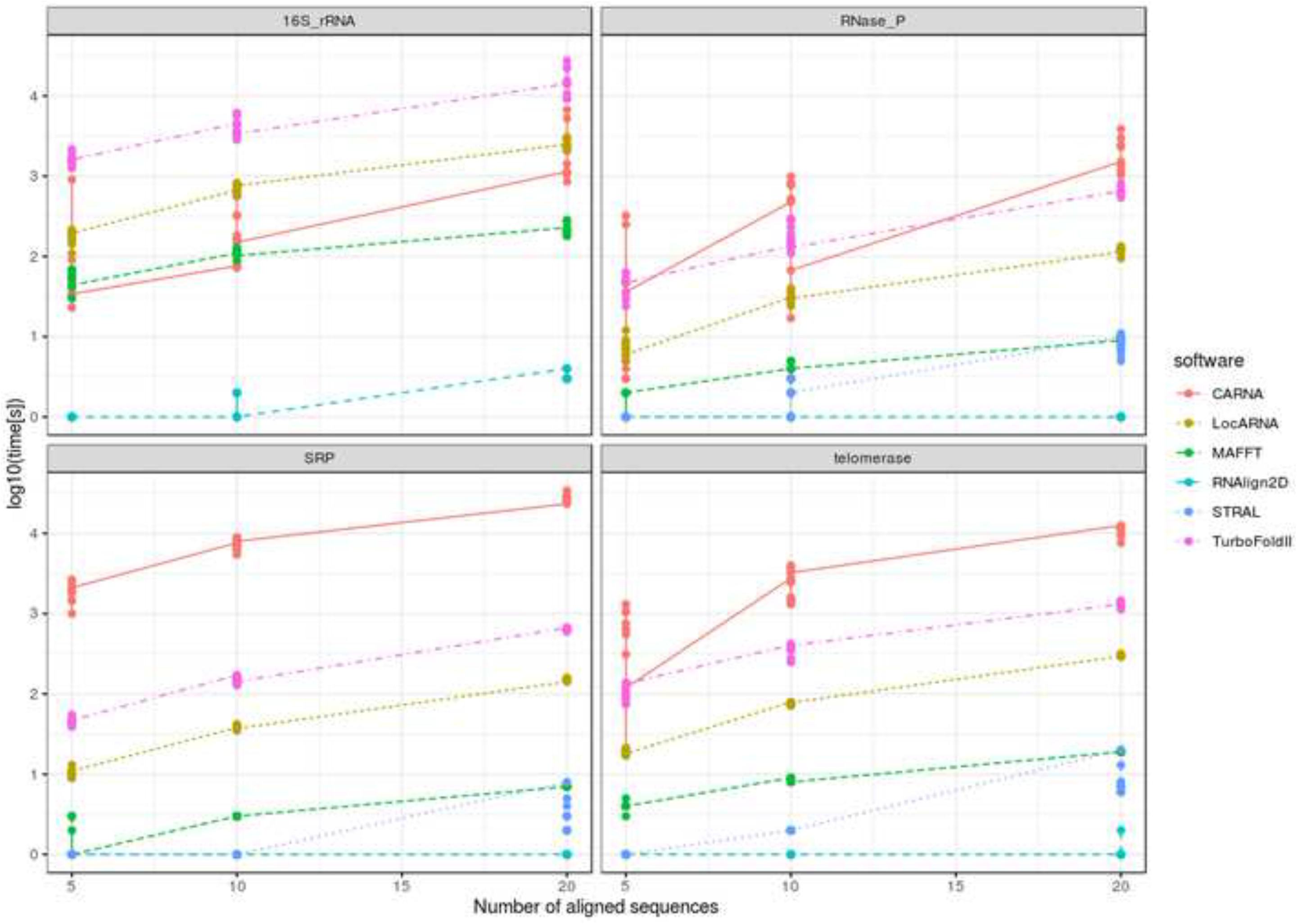
Comparison of alignment performance times between RNAlign2D, CARNA, LocARNA, MAFFT, STRAL, and TurboFold II for 10 sets of 5-, 10- and 20-sequences alignment from RNAStralign benchmark dataset. Measurement was not performed for STRAL and 16S rRNA dataset, because of occurring ‘segmentation fault’ error. Note that time [s] is shown at the log10 scale.

## Discussion

RNAlign2D is an extremely fast RNA alignment tool and thus allows the alignment of hundreds of RNA molecules in a very short time. It mediates alignment of RNA molecules with known 2D structures, where 2D structure is required as part of the input. RNAlign2D contains an option to model missing structures by using RNAfold from the ViennaRNA package [22], but in contrast to some existing programs (such as TurboFold II [5]), optimization of the structure prediction algorithm was beyond the scope of the project. Our tool is optimized for RNAs with known 2D structures. The biggest advantage of RNAlign2D is its faster speed in comparison to other tools, which was achieved by transformation of the sequence and 2D structure to pseudo-amino acid sequence followed by using a protein aligner (MUSCLE) to perform multiple sequence alignment (Figure 1). We chose MUSCLE aligner because of its good performance between 200 and 1000 sequences, which in our opinion would be the most common range of sequence number for RNAlign2D [23]. It is worth noting that the pseudo-amino acid term introduced in this paper refers to the method of encoding RNA sequence and 2D structure information as amino acid sequence, although it shares no similarities with pseudo amino acid composition (PseAAC) introduced by Chou, 2001 [24].

Overall, the RNAlign2D alignment performance (as indicated by SPSs and PPVs) is similar to LocARNA, CARNA, and TurboFold II, but RNAlign2D aligned the RNA sequences several hundred times faster than those tools. In some cases (e.g. RNase P and telomerase), it produced better alignment. In comparison to MAFFT and STRAL, RNAlign2D produced better alignment in the majority of benchmark datasets. However, alignment accuracy was strongly dependent on the RNA family and the different average pairwise sequence identity (APSI) values of the aligned sequences. Based on our benchmark results, RNAlign2D can be recommended as a first-choice tool for the alignment of large numbers of sequences with an APSI ≥ 50%. For instance, it can be used to align all members of a particular RNA family or all known tRNA isoacceptors/isodecoders for a specific amino acid. Results of such alignments can be further utilized to perform and/or improve 3D structure modeling.

For sequences with a low APSI (e.g. SRP RNA sequences in the RNAStralign benchmark, with average APSI = 38.7%), the performance of alignment with RNAlign2D was worse than that with LocARNA, CARNA, TurboFold II and MAFFT. It can be expected that a scoring matrix optimized for multiple RNA families could be sub-optimal for at least some of these families, including SRP in this case. We observed that in comparison to the SRP reference alignments, RNAlign2D introduced in general fewer gaps, especially in the stem regions and single-nucleotide bulges. Additionally, the introduced gaps are usually longer. This issue can be solved by changing the parameters in the scoring matrix, decreasing gap-opening penalty, or creating a scoring matrix optimized for the particular RNA family.

In terms of structural alignment quality, measured as mean structural distance between consensus structure and all structures in the input, RNAlign2D outperforms tools that use RNA structure prediction (STRAL and TurboFold II), which was expected. In comparison to other tools that utilize known RNA structure (LocARNA and CARNA), our tool was worse in the cases of telomerase and SRP, and at a very similar level for other datasets. It is worth noting here that better sequence alignment does not always mean smaller structural distance (as for the telomerase dataset).

We believe that there is still field for improvement of our approach in the future. To perform the best benchmark possible, we decided to use most of the available alignments for benchmark purposes. Therefore our training set was very limited. In case of the more manually curated structural alignments were available, it might be possible to introduce machine learning methods for optimization of either parameters specified in this publication or even each of the scoring matrix parameters.

## Conclusions

In conclusion, RNAlign2D uses a novel approach to align RNAs with known 2D structures, and with the growing number of experimentally determined RNA 2D structures, this approach will be further improved by optimization of scoring matrices for the particular RNA families and/or utilizing different aligners. It offers a reliable compromise between the computationally demanding approaches and fast, but much less accurate ones.

## Materials and Methods

### Benchmark – sum-of-pair-scores (SPSs) and positive predictive values (PPVs)

For benchmark purposes, RNAlign2D was compared with LocARNA (version 1.9.2.3) [7] and CARNA (version 1.3.4) [8], which represent other tools that use a fixed 2D structure for multiple RNA alignment, but also TurboFold II (version 6.2) [5] and MAFFT (version 2) [6], which produce the best alignments in another benchmark [5], and STRAL (version 0.5.4) [17] (with ViennaRNA 1.8.5 [25]), which uses a similar approach to encode sequence and structure. We used two available benchmark datasets: BraliBase 2.1 (k2, k3, k5, k10 and k15, where k indicates the number of aligned sequences) [20] and the dataset in RNAStralign [5]. First, we excluded tRNA sequences from BraliBase 2.1 to avoid a bias towards sequences whose identities are in the ‘twilight zone’ and range from 40% to 60%, most of which are tRNAs [5]. The BraliBase 2.1 dataset does not contain information about the secondary structures of aligned RNA molecules. Therefore, we first downloaded data indicating the secondary structures of all RNAs in the RFAM database [26], which was used to create the BraliBase 2.1 benchmark dataset, from the bpRNA-1m database [3]. Next, we converted the downloaded .ct files to dot-bracket format. To that end, we first removed all commentary lines from the .ct files using a custom Python script and then performed format conversion with the ct2dot tool from the RNAstructure package [27]. Finally, we used a custom Python script to add 2D structures to the BraliBase 2.1 raw.fa files and saved only the files that contained 2D structures for all sequences. Additionally, for files used as input for LocARNA and CARNA, we added ‘ #FS’ (which is required to align fixed 2D structures) to the end of each 2D structure line. For MAFFT, STRAL, and TurboFold II, we used regular fasta files containing only sequence as input. A complete list of files used, together with overlapping with k7 dataset used for optimization of the scoring matrix, is provided in Supplementary Table S3.

The benchmark on RNAStralign dataset was made as described by Tan et al. [5]. Namely, we generated 200 groups of 5, 10 or 20 sequence homologs selected from 16S rRNA sequences from Alphaproteobacteria, RNase P RNA sequences (bacterial type A subfamily), signal recognition particle (SRP) RNA sequences (protozoan subfamily), and telomerase RNA sequences.

In the case of 16S rRNA sequences from Alphaproteobacteria, we observed differences between some sequences in the ct files used as a test set and fasta file with reference alignment. Therefore, we first removed the sequences that differed from both the test and reference sets (RNAStralign IDs AB242948, AF301221, AY306224, AY436803, AY466761, AY785314, D14426, D14427, D14428, D14429, D14430, D14434, D14435, D84526, DQ303351, M803809, U71005, X79735, and X79738) and then proceeded to selection and analysis.

Sequences from the protozoan SRP reference alignment file contain a considerably higher number of unknown bases (Ns) than the same sequences in the test dataset used to perform alignments. Therefore, we utilized a custom Python script to replace unknown bases in the reference sequences based on the test dataset sequences and then employed these corrected reference sequences to calculate alignment accuracy.

We ran LocARNA, CARNA, STRAL, TurboFold II, and RNAlign2D (‘simple’ mode) with the following default parameters to align the complete benchmark datasets: #locARNA, mlocarna $file.raw.fa; #CARNA, mlocarna –pw-aligner carna $file.raw.fa; #STRAL, ./stral $file.fa; TurboFold II, ./TurboFold $file.config.txt (Mode = MEA, Gamma = 0.3, Iterations = 3, MaximumPairingDistance = 0, Temperature = 310.15); #RNAlign2D, rnalign2d -i $file.raw.fa -o$file.raw.fa.out. MAFFT was used in mxscarna mode, to predict RNA 2D structure # ./mafft_mxscarnamode $file.fa.

In the next step, SPSs and PPVs were calculated for each alignment. The output files of LocARNA and CARNA are in ClustalW aln format. To perform the calculations, we converted these files to FASTA format using the fasconvert tool from the FAST package (version 1.06) [28]. The output of RNAlign2D is a modified FASTA format including a header followed by a line containing the sequence and 2D structure in dot-bracket format. Therefore, the 2D structure line was removed using sed (sed ‘n; n; d’ < $file.raw.fa.out > $file.out.fasta). Other programs used in benchmark return output in fasta format, but STRAL put the empty line between aligned sequences. This empty line was removed using sed (sed -i ‘/^$/d’ $file.fa.out). FASTA files were sorted using a custom Perl script. SPS values were calculated using the compalignp program [29], where they are defined as the averaged identity over all N(N-1)/2 pairwise alignments. PPVs were calculated by applying a modified Python script used by another group [5]. Firstly, positions for each nucleotide in the test set and real set were calculated. In the next step, columns for each position were generated. Then the common part between columns (true positives) and difference between the test set and real set (false positives) were calculated. PPV was defined as the ratio of true positives to the sum of true positives and false positives.

To compare the mean SPSs and PPVs from RNAlign2D and other benchmarked programs, we applied the two-sided t-test, because of its better performance in comparison to non-parametric statistical test for large sample sizes, also when analyzed data are not normally distributed [30,31].

### Structural distance

To compare structural alignment accuracy between benchmarked programs, we calculated a mean from structural distances between consensus structure from each alignment and every single structure taken as input to the alignment, using RNAdistance (string alignment and full distance) from ViennaRNA package [22]. Consensus structures were calculated using custom Python script. We were unable to retrieve secondary structures predicted by MAFFT, therefore we excluded MAFFT from this analysis. t-test was used to measure statistical significance between mean structural distances. For the scoring matrix optimization purposes on k7 BraliBase 2.1 dataset 1 – (mean_distance/length of consensus structure) was used as a structural distance score.

### Alignment time

To determine the time required to perform each alignment, we used 40 groups of 5, 10 or 20 sequence homologs from the RNAStralign benchmark dataset. The LocARNA, CARNA, TurboFold II, MAFFT, STRAL, and RNAlign2D running times for each group were measured using the bash ‘time’ command.

### Figures

Figures 1–5 and 7–9 were generated using ggpubr package [32] with R.3.6.3 [33].

## Availability and Requirements

Project name: RNAlign2D

Project home page: https://github.com/tomaszwozniakihg/rnalign2d

Operating system(s): Linux, Mac OSX

Programming language: Python 3

Other requirements: MUSCLE (tested on version 3.8.31), pytest (tested on version 5.1.3), Vienna RNA (optional, tested on version 2.4.14)

License: MIT

Any restrictions to use by non-academics: no

## List of abbreviations

tRNA: transfer RNA
2D structure: secondary structure
rRNA: ribosomal RNA
SPS: Sum-of-pair score
PPV: Positive predictive value
RNase P: Ribonuclease P
SRP: Signal recognition particle
APSI: Average per sequence identity

## Declarations

Ethics approval and consent to participate: Not applicable

Consent for publication: Not applicable

## Availability of data and materials

All data generated or analyzed during this study are included in this published article and its supplementary information files.

## Authors’ contributions

Conceptualization, T.W. and M.P.S.; Data curation, M.P.S.; Formal analysis, T.W., M.S., and M.P.S.; Investigation, T.W. and M.P.S.; Methodology, T.W., M.S., and M.P.S.; Resources, J.J.; Software, T.W.; Supervision, M.P.S.; Visualization, T.W., M.S., and M.P.S.; Writing – original draft, T.W. and M.P.S; Writing – review & editing, T.W., M.S., J.J., and M.P.S.

All authors have read and approved the final version of the manuscript.

## Acknowledgements

We thank Dr. David Mathews and Dr. Zen Tan for providing useful scripts to calculate alignment accuracy, Dr. Tomasz Górecki for discussions of statistical analysis, Matisa Alla, Amber Baldwin and Kimberly Wellman for critical reading the manuscript.

Some part of this work was previously presented as a poster entitled “RNAlign2D- RNA sequence and structure multiple alignment tool, based on pseudo-amino acids substitution matrix”, at the Autumn Workshop PTBI 2020.

## Figures, tables, and additional files

Supplementary Figure 1. (A) Structure conversion to a pseudo-amino acid sequence for RNA with higher-level pseudoknots. (B) Conversion of structure elements to pseudo-amino acids and their scores (left) and the default scoring matrix (right).

Supplementary Table 1. Mean sum-of-pair scores (SPS) and positive predictive values (PPVs) with standard deviations obtained in BraliBase2.1 and RNAStralign benchmarks. In the highlighted fields, values differed between the full BraliBase2.1 benchmark (top values) and a smaller version of benchmark, where datasets containing ≥ 2 (k2, k3), ≥ 3 (k5), and ≥ 5 (k10) common sequences with k7 dataset were excluded (bottom values in parentheses).

Supplementary Table 2. Running time measurement for RNAlign2D in comparison to other aligners.

Supplementary Table 3. Bralibase2.1 dataset used to prepare benchmark. Additional sheet contains the numbers of overlapping sequences between the k7 dataset used for scoring matrix optimization and other Bralibase2.1 datasets.

