## Supplementary figures and images for "RNAlign2D – a rapid method for combined RNA structure and sequence-based alignment using a pseudo-amino acid substitution matrix"

### Supplementary Figure 1

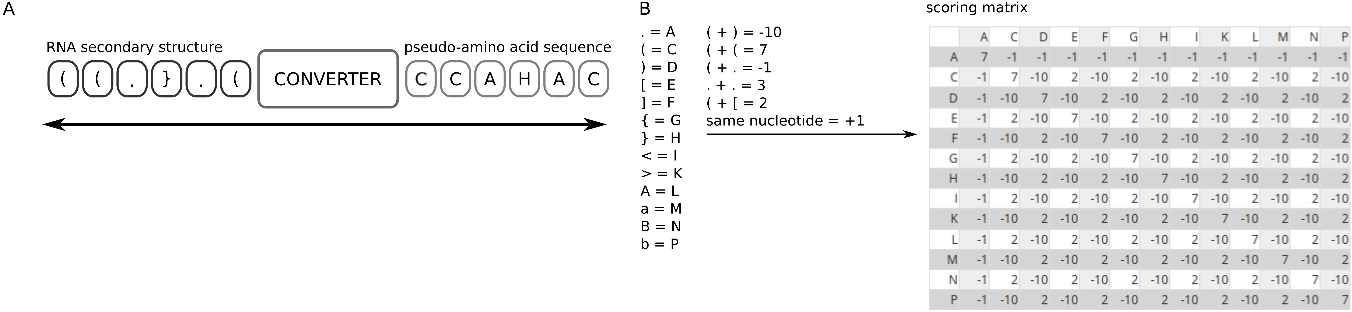
